# Infection dynamics in the urinary tract - The importance of non-planktonic phases during early infection and resistance evolution

**DOI:** 10.1101/2023.10.23.563535

**Authors:** Michael Raatz, Karolina Drabik, Amanda de Azevedo-Lopes, Arne Traulsen, Bartłomiej Wacław

**Affiliations:** Department of Theoretical Biology, Max Planck Institute for Evolutionary Biology, Plön, Germany; Dioscuri Centre for Physics and Chemistry of Bacteria, Institute of Physical Chemistry (ICHF), Polish Academy of Sciences, Warsaw, Poland

**Keywords:** Resistance evolution, evolution-informed medicine, antibiotic therapy, mathematical modelling

## Abstract

Treatment of urinary tract infections (UTIs) and the prevention of their recurrence is a pressing global health problem. During the infection establishment, pathogenic bacteria may attach to and invade into the bladder tissue, and thus access non-planktonic phases besides the bladder lumen. Planktonic, attached, and intracellular bacteria face different selection pressures from physiological responses (frequent micturition and immune responses), or antibiotic treatment. Acknowledging this heterogeneity of selection pressures is essential to increase treatment efficacy, reduce resistance evolution and develop new treatments. Here, we present a mathematical model capturing the initial infection phase to unravel the effects of this heterogeneity on the ecological and evolutionary dynamics of UTIs. We explicitly represent planktonic bacteria in the bladder lumen, bacteria attached to epithelial cells of the bladder wall, and bacteria that have invaded these epithelial cells. We find that access to non-planktonic compartments increases the risk of infection establishment substantially. With antibiotic treatment, the trajectories of resistance evolution are shaped by the accessibility of the intracellular compartment and antibiotic permeation into the epithelial cells. Further, we evaluate conditions where probiotic pretreatment can reduce the pathogen load during an infection and increase the efficacy of accompanying antibiotic therapies. We find that faster-growing probiotic strains achieve a stronger increase in antibiotic treatment efficacy, although too strong antibiotic treatment can diminish this effect. Our study shows that accounting for the selection heterogeneity is crucial to understand the eco-evolutionary dynamics of UTIs, and that mathematical modeling can help to steer this complex system into a healthy state.

**Competing Interests:** The authors declare no competing interests.

## Introduction

Urinary tract infections (UTIs) are a major health problem that affects millions of patients globally every year. Being one of the most common bacterial infections, and often recurrent, UTIs present a considerable risk of antibiotic resistance evolution (Flores-Mireles et al., 2015; Stracy et al., 2022), and contribute substantially to this global health burden (Murray et al., 2022). A large body of research has investigated the course of UTIs both experimentally and clinically to elucidate the infection dynamics as well as the host defense mechanism and the impact of treatment (see e.g. Cox and Hinman, 1961; Sobel, 1997; Mulvey et al., 2000; Justice et al., 2004; Walters et al., 2012; Murray et al., 2021). Despite these efforts, recurrent infections are common and the problem of evolution of resistance against antibiotics is not resolved, often even resulting in multidrug resistance (Flores-Mireles et al., 2015; Murray et al., 2021). A better understanding and novel treatment approaches, such as immunotherapies or probiotic treatment, are currently investigated to overcome these prevailing limitations (Li et al., 2022; Sihra et al., 2018; Kenneally et al., 2022).

The urinary tract consists of multiple distinct spatial environments for uropathogenic bacteria. Accounting for these heterogeneous environments is crucial to understand the course of infections. In the lumen of the bladder, regular bladder voiding results in frequent clearance, but incomplete voiding is associated with higher infection risks. Consequently, urological voiding restrictions (incontinence, presence of a cystocele, postvoiding residual urine) present strong associations with recurrent UTIs (Raz et al., 2000). The frequent voiding motivated the classical assumption of a sterile bladder as the healthy state, which has now been revoked (Ackerman and Chai, 2019). The bladder epithelium consists of umbrella cells to which bacteria from the bladder microbiome but also pathogens can adhere. Aided by adhesins and pilii, bacteria transition from the planktonic phase into this attached phase from where they may invade into the first layer of epithelial cells in the bladder tissue (Schwartz et al., 2011). Here, the pathogens form biofilm-like intracellular bacterial communities (IBCs) which provide opportunities for growth and protection from immune responses (Anderson et al., 2003).

These three compartments of planktonic, attached and intracellular phase generate a spatial hetero-geneity of differing ecological and evolutionary selection pressures. Bladder voiding is able to remove most of the bacteria in the lumen of the bladder, causing a major population size bottleneck, and is understood as a physical protection mechanism against urinary infections (Cox and Hinman, 1961). However, its effect on attached and intracellular bacteria is likely much smaller. Immune responses, e.g. secretion of antibodies, recruitment of neutrophils, and exfoliation driven by mast cells (Mulvey et al., 2000; Choi et al., 2016), in turn, will have the strongest effect on the attached and intracellular bacteria. Further, the biofilm-like phase of intracellular growth slows down growth and leads to physiological changes such as a coccoid shape (Sharma et al., 2021). If treated with antibiotics, the permeation of the drug as well as the biophysical properties of the intracellular biofilm-like structure limit the drug exposure of the intracellular bacteria (Singh et al., 2010), while planktonic and attached bacteria are exposed to higher drug concentrations (Trubenová et al., 2022b). Such spatial hetero-geneity of the antibiotic selection pressure may facilitate resistance evolution in bacteria (Greulich et al., 2012; Hermsen et al., 2012; Moreno-Gamez et al., 2015; Nicholson and Antal, 2019; Trubenová et al., 2022a). To more efficiently cure UTIs, prevent recurrent UTIs, and limit resistance evolution a mechanistic understanding of the dynamics during early infection is imperative.

In this study, we incorporate both ecological and evolutionary selection pressures into a mathematical model of bacterial population dynamics in the three compartments of planktonic, attached and intracellular growth phase of uropathogenic bacteria to investigate the effect of this heterogeneity on the infection dynamics. First, we study the effect of general accessibility of the attached and intracellular compartment to the invading bacteria. Then, using a variant of the model in which both compartments can be colonised by the bacteria, we explore the effect of the invasion rate at which bacteria can internalize into the epithelial cells from the attached phase in relation to the permeation of an antibiotic drug into the epithelial cells. Finally, we investigate when colonization resistance by probiotic prophylaxis can limit the spread of a pathogenic infection.

## Methods

We construct a mathematical model to represent the initial dynamics of urinary tract infections within the lumen of the bladder, on the surface, and inside of the outermost layer of epithelial cells of the bladder wall. Motivated by the differential selection pressures acting on bacteria in the planktonic, attached, and intracellular phases we define a distinct compartment for each of these three phases. To account for the inherent between-cell heterogeneity of epithelial cells, we further split the intracellular compartment into independent subpopulations for each epithelial cell, thus representing the infection dynamics of individual epithelial cells. Assuming that the bladder surface does not have any large-scale heterogeneities, we model 1 mm^2^ of the bladder, i.e. we consider a system size of 1 mm^2^ of bladder surface and the volume of 1 mm^3^ on top of this surface. We place *N* = 250 epithelial cells on this surface, thus assuming an average area per epithelial cell of 4000 µm^2^. Table 1 provides a summary of all model parameters.

**Table 1.** Reference parameter set. Deviations from these values are reported where applicable.

|  | Biological meaning | Value | Reference |
| --- | --- | --- | --- |
| $\beta_p, \beta_a$ | Maximum birth rate of extracellular bacteria | $1.19 \text{ h}^{-1}$ | (Forsyth et al., 2018)<sup>1</sup> |
| $\overline{\beta_{ci}}$ | Average maximum birth rate of intracellular bacteria | $b/2 = 0.595 \text{ h}^{-1}$ | (Sharma et al., 2021)<sup>2</sup> |
| $\delta_p, \delta_a$ | Death rate of extracellular bacteria | $b/100 = 0.0119 \text{ h}^{-1}$ | this work |
| $\overline{\delta_{ci}}$ | Average death rate of intracellular bacteria | $b/100 = 0.0119 \text{ h}^{-1}$ | this work |
| $K_p$ | Carrying capacity for planktonic bacteria | $10^6 \text{ mm}^{-2}$ | (Forsyth et al., 2018) |
| $K_a$ | Carrying capacity for attached bacteria | $10^6 \text{ mm}^{-2}$ | this work<sup>3</sup> |
| $\overline{K_c}$ | Average carrying capacities for intracellular bacteria | 4000 | this work<sup>3</sup> |
| $\sigma_\beta$ | Standard deviation of intracellular max. birth rates | 0.089 25 | this work<sup>4</sup> |
| $\sigma_\delta$ | Standard deviation of intracellular death rates | 0.001 785 | this work<sup>4</sup> |
| $\sigma_K$ | Standard deviation of intracellular carrying capacity | 600 | this work<sup>4</sup> |
| $\lambda_a$ | Attachment rate of planktonic bacteria | $0.2 \text{ h}^{-1}$ | (Martinez et al., 2000)<sup>5</sup> |
| $\lambda_d$ | Detachment rate of attached bacteria | $0.005 \text{ h}^{-1}$ | (Seiler et al., 2017) |
| $\nu$ | Invasion rate of attached bacteria | varied, $10^{-5} \dots 10^{-2} \text{ h}^{-1}$ | |
| $\mu$ | mutation rate per birth | $5 \times 10^{-8}$ | (Komp Lindgren et al., 2003) |
| $\kappa$ | Antibiotic kill-fraction | 0.999 | (Hofsteenge et al., 2013)<sup>6</sup> |
| $\phi$ | Intracellular permeation of treatment | varied, $0 \dots 1$ | |

<sup>1</sup> Corresponding to a doubling time of 35 min.
<sup>2</sup> Corresponding to a doubling time of 70 min.
<sup>3</sup> Set such that the total carrying capacities in all three compartments are equal. See further reasoning in the main text.
<sup>4</sup> Set to ensure equal coefficients of variation of 0.15 for intracellular birth and death rate and carrying capacity.
<sup>5</sup> Martinez et al. (2000) show adherence of 46% of inoculated bacteria after two hours.
<sup>6</sup> Corresponding to persister fractions of $10^{-3}$ as observed in (Hofsteenge et al., 2013).

Bacteria can migrate between compartments, replicate, potentially creating mutant offspring lineages, and die. We assume logistic density-dependent birth rates and constant death rates in all compartments. The birth rates *b_i_* and death rates *d_i_* in compartment *i* thus take the form

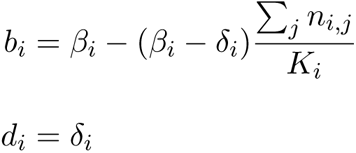

with maximum birth rate *β_i_*, intrinsic death rate *δ_i_*, and carrying capacity *K_i_*. Here, *n_i,j_* is the population size of lineage *j* in compartment *i* with *i ∈ {p, a, c*1*,…, cN}* for planktonic, attached and the *N* intracellular compartments. We denote the wildtype with *j* = 0 and the mutant lineages with *j* = 1, 2*,…*.

Further, we assume that planktonic and attached bacteria can replicate with the same maximum birth rate *β_p_* = *β_a_*, which we set to 1.19 h*^−^*^1^, arising from the assumption of exponential growth with a doubling time of 35 min (Forsyth et al., 2018). For the planktonic phase, we assume a carrying capacity for birth of *K_p_* = 10^6^ mm*^−^*^2^, which for a height of 1 mm corresponds to a typical bacterial volume density of 10^9^ ml (Forsyth et al., 2018). Assuming that at closest packing attached bacteria form a monolayer and require approximately 1 µm^2^ per cell, we set the carrying capacity for birth in the attached phase to *K_a_* = 10^6^ mm*^−^*^2^ and thus equal to the carrying capacity in the planktonic phase. Heterogeneity both in size and chemical properties among the epithelial cells likely leads to differing growth conditions for the intracellular bacterial subpopulations. Thus, we are assuming uncorrelated, normally-distributed birth rates, death rates, and carrying capacities for the subpopulations in each epithelial cell. Also, this heterogeneity prevents unrealistically synchronized infection events of the subpopulations which would lead to long-term oscillations in the number of infected epithelial cells. Mirroring the biofilm-like growth conditions within the epithelial cells, we assume that the average maximum birth rate of intracellular growth equals half of the extracellular maximum birth rate *β_p_*, resulting in an average doubling time of 70 min, in agreement with recent findings (Sharma et al., 2021). The average death rates are assumed to be equal in all compartments and correspond to 1*/*100 of the planktonic maximum birth rate. As average carrying capacity for intracellular bacteria birth we assume 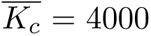 bacteria per cell as it ensures that the total intracellular carrying capacity equals the planktonic and attached carrying capacities. This choice corresponds to a maximum intracellular bacterial volume occupation of 25% when assuming an average height of epithelial cells of 4 µm and an average bacterial volume of 1 µm^3^.

Consequently, we assume

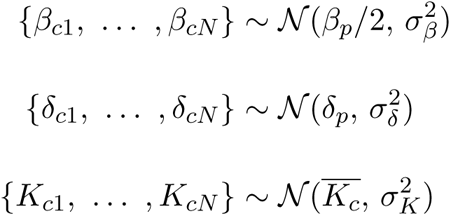

for the parameters determining the growth conditions of intracellular populations. Upon birth, we assume a probability of resistance mutation of *µ* = 5 *×* 10*^−^*^8^ (Komp Lindgren et al., 2003). We assume that in the absence of antibiotics these mutations are neutral. Each newly created mutant founds a lineage that we track in parallel to the wildtype lineage and all other lineages. Migration between the planktonic and attached phases is determined by the attachment rate *λ_a_* and detachment rate *λ_d_*. Attached bacteria can invade an epithelial cell at rate *ν*. We assume that all epithelial cells have the same probability of being invaded. As estimating this invasion rate is difficult, we will consider a wide range of possible values for this parameter. Once the subpopulation of intracellular bacteria in an epithelial cell reaches 95% of the epithelial cell’s carrying capacity all bacteria from that subpopulation are released to the planktonic phase, mimicking a burst event similar to pyroptosis or exfoliation (Justice et al., 2004; Choi et al., 2016; Wu et al., 2019). After such a burst event, the burst epithelial cell is replaced with an empty cell with identical parameters *β_ci_*, *δ_ci_* and *K_ci_*.

### Perturbations

Bladder voiding exerts a repeated perturbation on the bacteria in the planktonic phase. Given the typical voiding frequency in humans of 6 d*^−^*^1^ to 8 d*^−^*^1^ (Lukacz et al., 2009), we assume that every four hours 90% of the planktonic bacteria are removed on average (Maramba et al., 2022). Bacteria in other compartments remain unaffected. Antibiotic treatment is modelled as a repeated population bottleneck that removes almost all (*κ* = 99.9%, Hofsteenge et al. (2013)) wildtype bacteria from the planktonic and attached compartments. The membrane permeability to the antibiotic drug, i.e. the drug permeation *ϕ*, determines whether treatment also acts within the epithelial cells and thus defines the difference in selection for resistance between the extracellular and intracellular compartments. The fraction of bacteria that is removed by one treatment cycle from the intracellular compartment thus equals *ϕ κ*. Therefore, lower drug permeation can create a refuge for the bacteria. The bacterial invasion rate *ν* determines whether this potential refuge can be accessed by the bacteria. Accordingly, we will investigate the effect of these two parameters on the evolution of resistance for different frequencies of antibiotic treatment.

**Figure 1.**
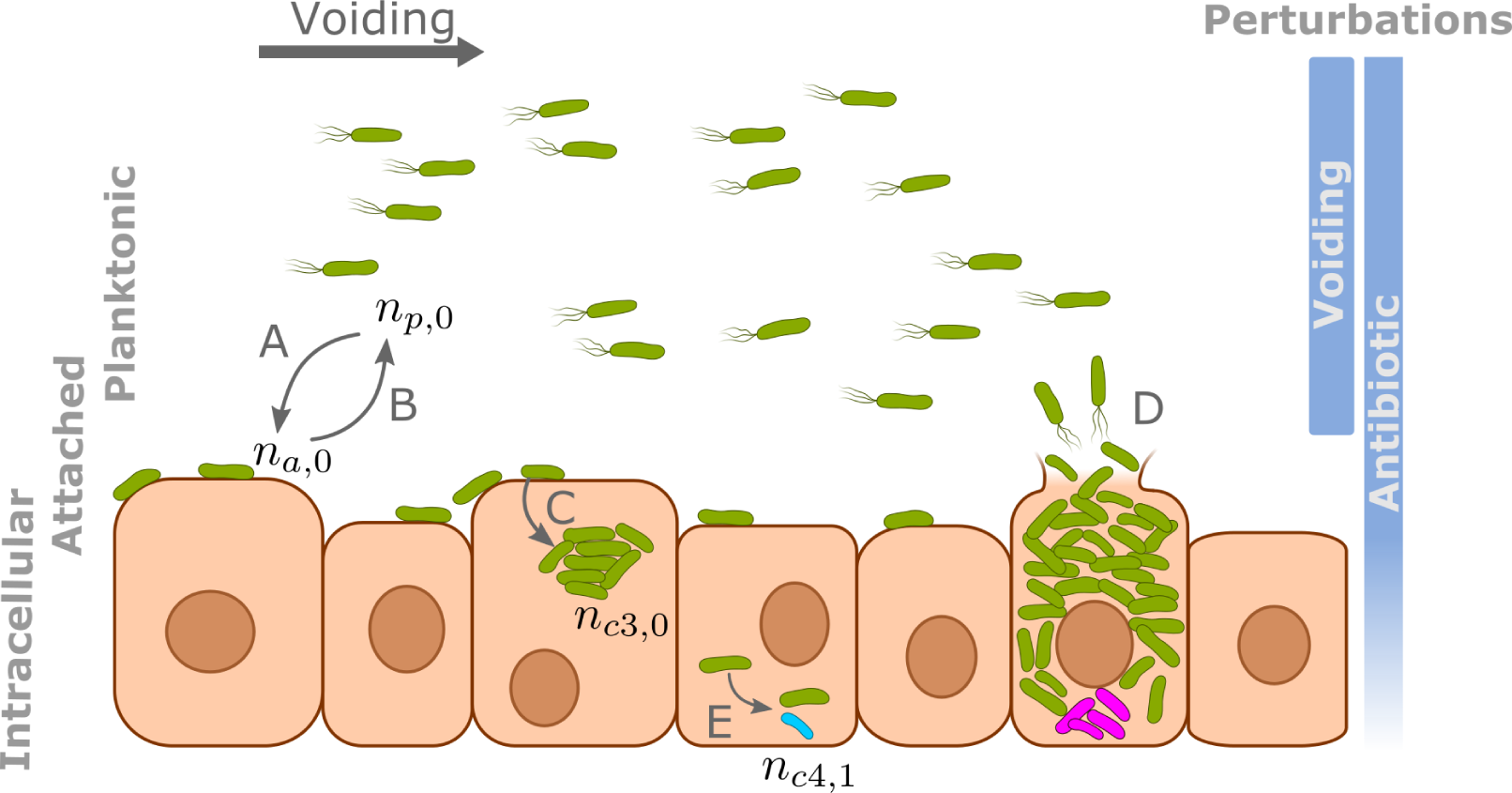
Sketch of the mathematical model for urinary tract infections. We are considering the three compartments of planktonic, attached, and intracellular bacteria. Besides birth and death, the model includes the attachment (A) and detachment (B) dynamics, the invasion into the epithelial cells (C), the bursting of epithelial cells once they become too crowded, resulting in the release of the intracellular population from that cell to the planktonic compartment, and the acquisition of resistance mutations (E). Resistant mutants result in new lineages that compete with other lineages in their compartment or epithelial cell. We consider the two perturbation types of bladder voiding and antibiotic treatment. Voiding targets only the planktonic compartment. Antibiotic treatment may target all compartments, but intracellular killing rate depends on the treatment permeation *ϕ*.

### Stochastic simulations

We implement the above model as a discrete time stochastic simulation in Python (version 3.10). The pseudocode in Supplementary Box 1 illustrates the simulation workflow. We inocculate the computational model system with 10^4^ planktonic cells and iterate through time with a step size of dt = 0.01h, tracking the abundances of all lineages in all compartments. The simulation code is available on Zenodo at https://doi.org/10.5281/zenodo.10025283. The data files have been deposited at https://doi.org/10.5281/zenodo.10025291.

## Results

Tracking the spread of an infection in space and time, we can understand the selection pressures acting on the pathogen population during the initial phases of urinary tract infections. We first show the importance of spatial heterogeneity by considering the effect of bacteria migrating between the three compartments (planktonic, attached, and intracellular). Then, we explore the interaction of ecological and evolutionary dynamics in this system in more detail by considering a clinically relevant problem of the evolution of antibiotic resistance. Finally, we will explore whether prophylactic pretreatment with a less invasive strain, i.e. a probiotic, can prevent pathogenic infections and limit the spread of resistance mutants.

### Importance of non-planktonic compartments

Voiding, even when assuming an incomplete reduction of the planktonic population, exerts a considerable perturbation and can achieve protection against infection, particularly if the pathogenic bacteria do not have access to the attached or intracellular compartments (Fig. 2). Supplementing this natural defense mechanism with antibiotic treatment further decreases the risk of infection establishment. Similarly, shorter voiding and treatment intervals lead to a lower risk of infection establishment. Access to the attached and intracellular compartments, however, obstructs the protective effect of voiding and necessitates antibiotic treatment at a treatment interval that is small enough to prevent regrowth between treatment applications (marked by the horizontal black line in Fig. 2a). The treatment period necessary for prevention of infection establishment decreases even further if the pathogenic bacteria can access the intracellular compartment and if treatment permeation is imperfect (*ϕ <* 1), thus resulting in an efficient shelter from both voiding and antibiotic treatment.

**Figure 2.**
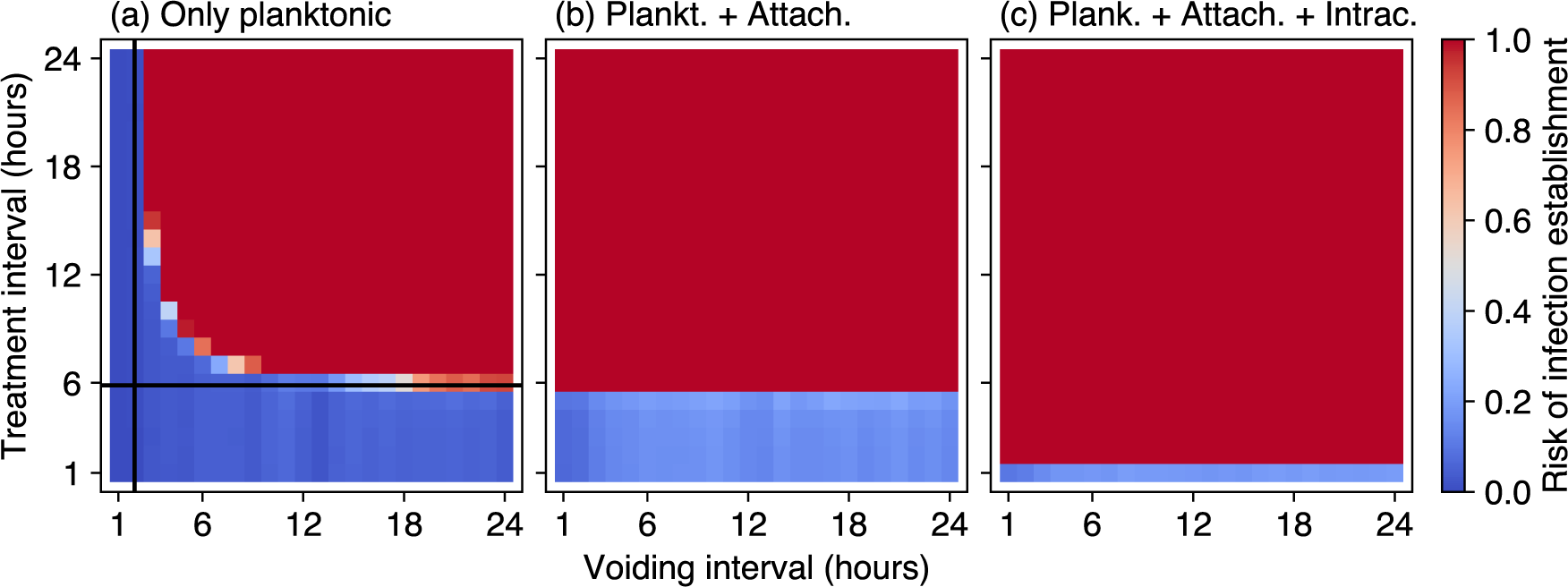
A model with only the planktonic compartment underestimates the establishment probability of urinary tract infections. Establishment of an infection may be prevented by a combination of bladder voiding and antibiotic treatment, but depends on their temporal intervals as well as on the accessibility of the spatial compartments to bacteria (panel (a) - only planktonic, panel (b) - planktonic and attached compartments, panel (c) - planktonic, attached and intracellular compartments accessible). The risk of infection establishment is measured as the proportion of 1000 replicates with non-extinct bacterial populations at *t* = 100 h after an initial infection event at *t* = 0 h with a planktonic inocculum of 10^4^ bacteria. First voiding occurs at *t* = 4 h and is repeated regularly at the voiding interval (horizontal axis). Treatment is first applied at *t* = 30 h and is repeated regularly at the treatment interval (vertical axis) with a permeation of *ϕ* = 0.75, i.e. a slightly reduced intracellular treatment effect. The black lines in panel (a) mark the voiding (vertical) and treatment (horizontal) interval that would minimally be required to prevent the spread of a wildtype population of pathogenic bacteria that grows exponentially at rate *β_p_ −δ_p_*. These lines thus indicate the threshold intervals that would be sufficient to clear the infection in the absence of spatial heterogeneity or resistance evolution. In the simulations, access to the attached and intracellular compartments is restricted by setting the attachment rate *λ_a_* or the invasion rate *ν* to zero, respectively.

### Eco-evolutionary dynamics of urinary tract infections

In the following, we will consider the scenario where bacteria can access all three spatial compartments and investigate the ecological and evolutionary dynamics in more detail. If left untreated, bacteria can colonize all available compartments of the model, i.e. the planktonic, the attached, and the intracellular one (Fig. 3a). Right after inocculation, we see a period of low planktonic densities due to the losses from voiding. Some of the planktonic bacteria colonize the epithelial cell surface which provides shelter from voiding. After colonization of the surface, the first bacteria invade the epithelial cell layer and commence intracellular growth. The first invasion events into each epithelial cell occur quickly and almost all epithelial cells are colonized within hours after the initial infection. In these infected epithelial cells, bacteria grow and eventually cause the epithelial cells to burst and release all intracellular bacteria, which move to the planktonic compartment. From here, they are either quickly removed by voiding, or they escape voiding by attaching to the epithelial cell surface. This influx from attachment causes the attached population size to increase beyond the attached carrying capacity *K_a_*. The burst epithelial cells are replaced with empty epithelial cells leading to a drop in the fraction of infected epithelial cells before new invasion events again increase the fraction of infected epithelial cells. Mutations can occur in all compartments, but they are quickly lost in the absence of treatment. Adding antibiotic treatment changes these dynamics drastically (Fig. 3 panels b-d), endowing resistant mutants with a fitness benefit which helps them spread through all three compartments. As this spread is initially hindered by competition with wildtype bacteria, and mutants are initially rare, the planktonic and attached bacterial population sizes are smaller under treatment during the early phase of treatment. What happens next strongly depends on the treatment permeation *ϕ*, i.e. the intracellular effect of antibiotic treatment. For non-permeating treatment, the intracellular compartment provides a refuge to the wildtype population. In the absence of resistant mutants, this refuge maintains the wildtype population if treatment is not frequent enough, as shown in Fig. 4. In the presence of mutants, the remaining wildtype population delays the spread of resistant mutants via competition (Fig. 3b). If treatment is fully permeating the intracellular wildtype population is quickly removed and, in the absence of resistant mutants, the infection is cleared (Fig. 3d). In the presence of mutants, however, their take-over occurs rapidly due to their large fitness advantage. For intermediate permeation, however, we find that the intracellular populations are kept at intermediate levels for considerable times by the drug as bursting is prevented (Fig. 3c). Therefore, a partially permeating drug or a drug that is ineffective in the intracellular compartment due to other reasons maintains bacterial infections for potentially long periods of time. If a resistant mutant has evolved, it can however co-invade these epithelial cells that already harbour wildtype populations, outgrow the wildtype and cause the burst of the epithelial cells, and thus eventually clear the system from wildtype bacteria (Fig. 3c). These dynamics are mirrored in the average population sizes across all compartments, indicating a strong effect of both the invasion rate and the drug permeation if treatment is applied (Fig. 4). We find that a faster invasion rate *ν* increases the average population size as it provides better access to the intracellular compartment (Fig. 4a). Infrequent treatment leads to a moderate reduction in bacterial population sizes (Fig. 4b). For frequent treatment we find a strong reduction in average bacterial population sizes, particularly for either low or high drug permeation. As described above, intermediate permeation maintains an intermediate intracellular bacterial population size and thus allows for a larger average population size (Fig. 4c). Low permeation allows for intracellular growth, which leads to bursting and consequentially to the expulsion of intracellular populations from their refuge — and thus lower average intracellular population sizes. Therefore, the probability for clearing the infection is non-zero not only for frequent treatment with a high drug permeation, but interestingly also for low permeation and low invasion rate (Fig. 4 panels d-f). In this latter case, planktonic and attached bacterial subpopulations are suppressed by treatment and voiding. Invasion into the intracellular compartment is slow, and those intracellular subpopulations that have formed burst their epithelial hosts quickly, which transfers the bacteria to the planktonic compartment where they experience a high mortality.

**Figure 3.**
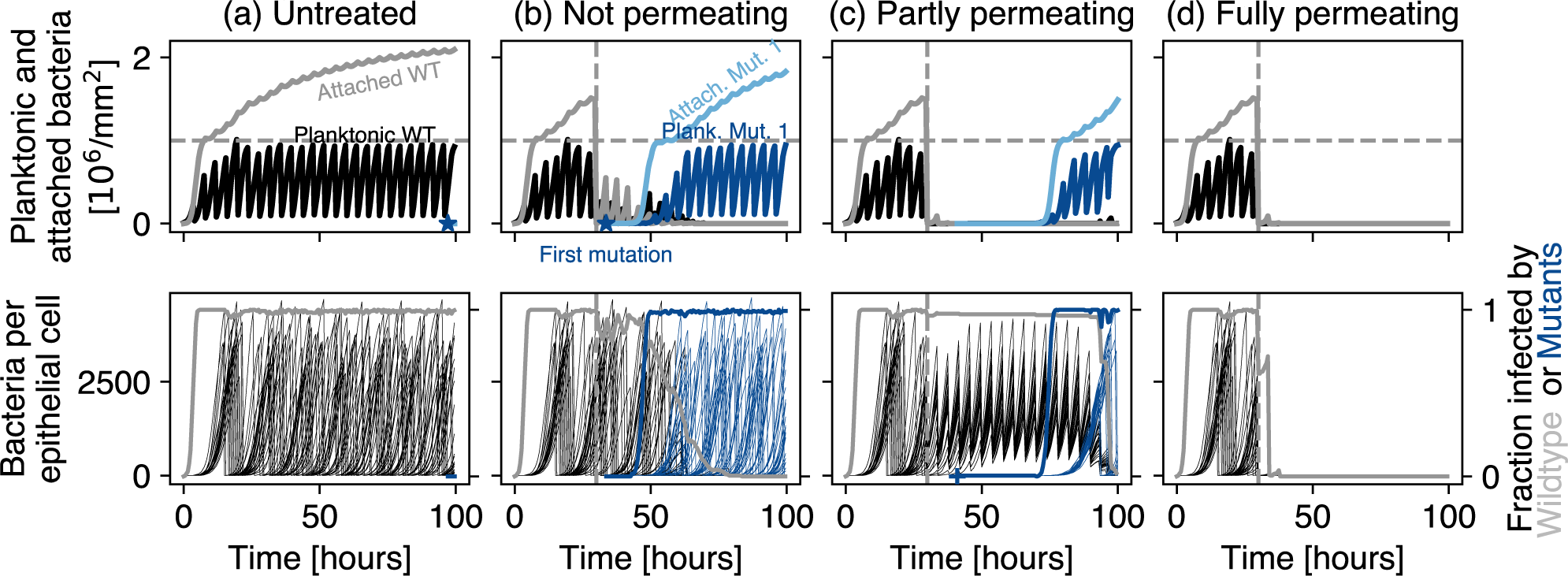
Exemplary population dynamics for different levels of treatment. Perturbations are: (a) voiding only, (b) voiding and frequent antibiotic treatment that does not permeate into the epithelial cells (*ϕ* = 0), (c) voiding and frequent antibiotic treatment that partially permeates into the epithelial cells (*ϕ* = 0.75), (d) voiding and frequent, fully permeating treatment (*ϕ* = 1). The invasion rate here is *ν* = 10*^−^*^3^ h*^−^*^1^. The top row shows the bacterial population densities of the planktonic and attached phases. The dashed horizontal line indicates the carrying capacities *K_p_* and *K_a_*. The bottom row represents the bacterial populations within each epithelial cell (thin lines, left vertical axis), as well as the fraction of infected epithelial cells (thick lines, right axis). If present, the vertical dashed line marks the first treatment event at *t* = 30 h, from whereon treatment is applied every 4 hours. For better representation only every 50^th^ intracellular population is shown. Symbols mark mutation events in the planktonic (circle), attached (star) and intracellular compartment (cross).

**Figure 4.**
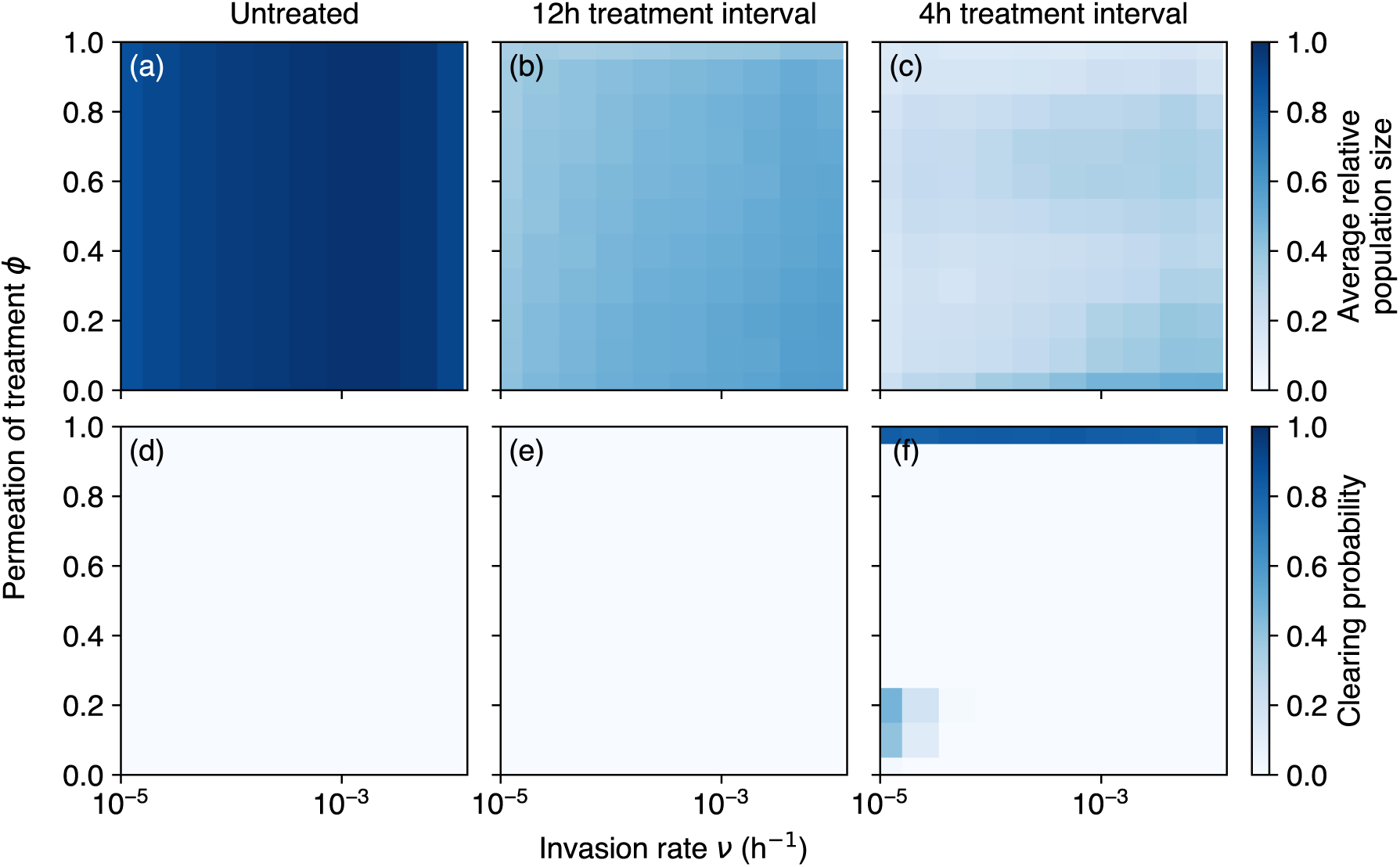
Effect of refuge accessibility and quality on average relative population size and clearing probability. Top row: Average population size for 1000 replicates between *t* = 80h and *t* = 100h in all compartments. Bottom row: Probability for clearing the infection for different perturbation levels. The perturbation levels are: voiding only (left column), voiding and infrequent antibiotic treatment (middle), and voiding and frequent treatment (right column). The axes represent the invasion rate *ν*, i.e. the rate at which an attached bacterium transitions into the intracellular compartment, and the permeation of treatment *ϕ*. Treatment starts from *t* = 30 h and is repeated every 12 hours in panels (b) and (e) and every 4 hours in panels (c) and (f). Population size is plotted relative to the total carrying capacity of the system 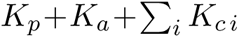 Clearing probability is measured as the proportion of 1000 replicates where the bacterial population is extinct at *t* = 100 h.

### Lineage tracing elucidates the origin of resistant mutants

Our stochastic simulations allow us to trace resistance evolution from the emergence of mutants to their eventual spread or extinction. The emergence of resistant mutants depends both on the presence of a large wildtype population and a high replication rate. Survival of these mutants and their eventual dominance over the wildtype requires a selection for resistance, such as the one imposed by antibiotic treatment. The three compartments (planktonic in the bladder lumen, attached to the epithelial cells, internalized in the epithelial cells) of our model represent the distinct ecological niches that bacteria face in UTIs and allow us to track their effect on the evolutionary trajectories during the evolution of antibiotic resistance.

We find that the treatment level strongly alters the origin of successful mutants, i.e. the compartment where a mutant lineage emerged that survives our simulation time (Fig. 5). An untreated bacterial population which only experiences voiding will acquire most of its successful resistant mutants in the planktonic phase, with some contributions from the intracellular phase for high invasion rates. When exposed to low-frequency antibiotic treatment, the attached phase gains much more importance as the origin of surviving resistant mutants. Instead, for high-frequency antibiotic treatment, we find a strong contribution of the intracellular phase to resistance emergence for intermediate permeation. These intricate patterns can be understood by a more detailed analysis of the population sizes and mutation rates in each compartment for each treatment level.

**Figure 5.**
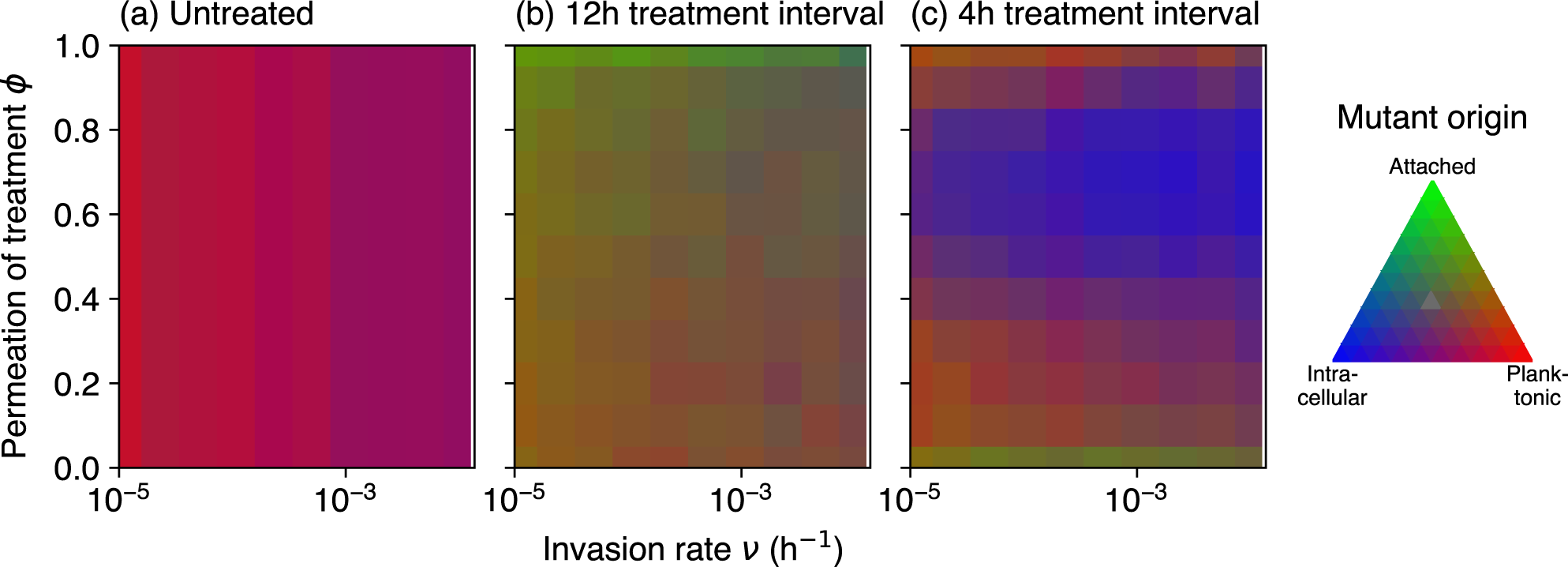
Origin of non-extinct mutants for the three treatment levels. Establishment of mutants from each compartment (planktonic, attached, intracellular) depends on the bacterial invasion rate *ν* and the treatment permeation *ϕ*, and is affected by the treatment scenario (same as in Fig. 4). The red, green and blue values for each pixel are assigned from the relative frequency of mutations in the three compartments, i.e. the average number of mutations in a specific compartment divided by the sum of all number of mutations averaged across 1000 replicates. Note that this figure does not represent absolute number of mutations.

The average population sizes presented in Fig. 4 are composed of wildtype and mutant subpopulations present across these three compartments (Fig. S1). In the planktonic compartment, the bacteria face frequent and complete population crashes by bladder voiding which keeps average planktonic population sizes low, but allows continued replication between voiding events because of competitive release. This replication fuels a large influx into the attached compartment, which, in the absence of antibiotic treatment, leads to population sizes beyond this compartment’s carrying capacity, thus prohibiting replication of the attached subpopulation (Fig. S1a). The attached compartment then supplies bacteria to the intracellular pool, at a rate proportional to the invasion rate *ν*. The larger wildtype subpopulations in the planktonic and internalized compartments for higher invasion rates result in a higher number of mutation events whereas the vanishing replication in the attached compartment generates few mutations (Fig. S2a). This leads to a positive correlation between average population size and the number of mutations only for the planktonic and intracellular compartments (Fig. S3). In the absence of antibiotic treatment, these mutants lack a selective advantage and are thus subject to stochastic extinction, indicated by minor average population sizes (Fig. S1a) and low survival probabilities during our observation period (Fig. S2a). Infrequent treatment, in combination with frequent voiding, strongly reduces population sizes in all compartments and shifts the balance towards the mutants, now giving them a selective advantage (Fig. S1b). Interestingly, this selective advantage depends on the treatment permeation parameter *ϕ*. For low permeation the wildtype still dominates in our observation window. Mirroring the population size pattern, most mutations in the planktonic compartment are observed for low permeation (Fig. S2b). Due to the frequent voiding, these mutants have a small chance of survival. The reduced population size in the attached compartment now allows for replication and a moderate number of mutations also in this compartment. Interestingly, the survival probability of these mutants is large and exceeds the survival probability of the mutants that originate from the planktonic and internalized compartments. More frequent treatment alters the balance between wildtype and mutant (Fig. S1c). Here, mutants reach high average population sizes in the attached compartment for low and high, but not for intermediate permeation, where wildtype cells dominate the intracellular compartment. As described above, these intermediate values of permeation, together with the high treatment frequency, prevent bursting of the epithelial cells without eradicating the intracellular populations. These sustained intracellular wildtype populations compete with the mutants and prevent their rapid take-over. Only after several rounds of treatment the mutants will eventually dominate due to their selective advantage and burst their epithelial host cell, thus clearing also the wildtype subpopulation. The sustained intracellular wildtype populations present replication hot-spots which give rise to many mutations whose survival probability, however, is often lower than the survival probability of mutants that arose in the attached compartment (Fig. S2c).

### Probiotic treatment to prevent infections

We have shown that the competition between wildtype and resistant mutants is an important determinant of infection dynamics and resistance evolution. In light of the risk of antibiotic treatment failure due to resistance evolution it is imperative that alternative or supplementary treatment types are developed against UTIs. Besides treating an acute infection, the prevention of the initial establishment of pathogenic bacteria in the urinary tract is an important aim to reduce the overall disease burden and increase the well-being of many patients affected by recurrent UTIs. One such approach is the prophylactic or adjuvant introduction of non-pathogenic bacteria into the urinary tract, i.e. probiotic pre-treatment, to prevent the establishment of a more pathogenic strain. In the following, we will explore whether less pathogenic bacterial strains that induce an asymptomatic bacteriuria (labeled ABU hereafter) could be used to prevent the spread of pathogenic strains (labeled PATH) via competition, and whether such probiotic treatment could also restrict the evolution of resistance to a parallel antibiotic treatment.

We assume that the increased pathogenicity of the pathogenic strain would manifest as a higher invasion rate *ν*, at the cost of a decreased birth rate relative to the probiotic strain. For the probiotic strain we thus assume an invasion rate at the lower end of the parameter range that was investigated above, but a high birth rate. Exploring this trade-off we find that pre-treatment with an ABU strain delays the spread of the pathogen (Fig. 6). This delay leads to a decreased pathogen population size (Fig. S4), most strongly if no antibiotic treatment is applied, and predominantly in the attached compartment.

**Figure 6.**
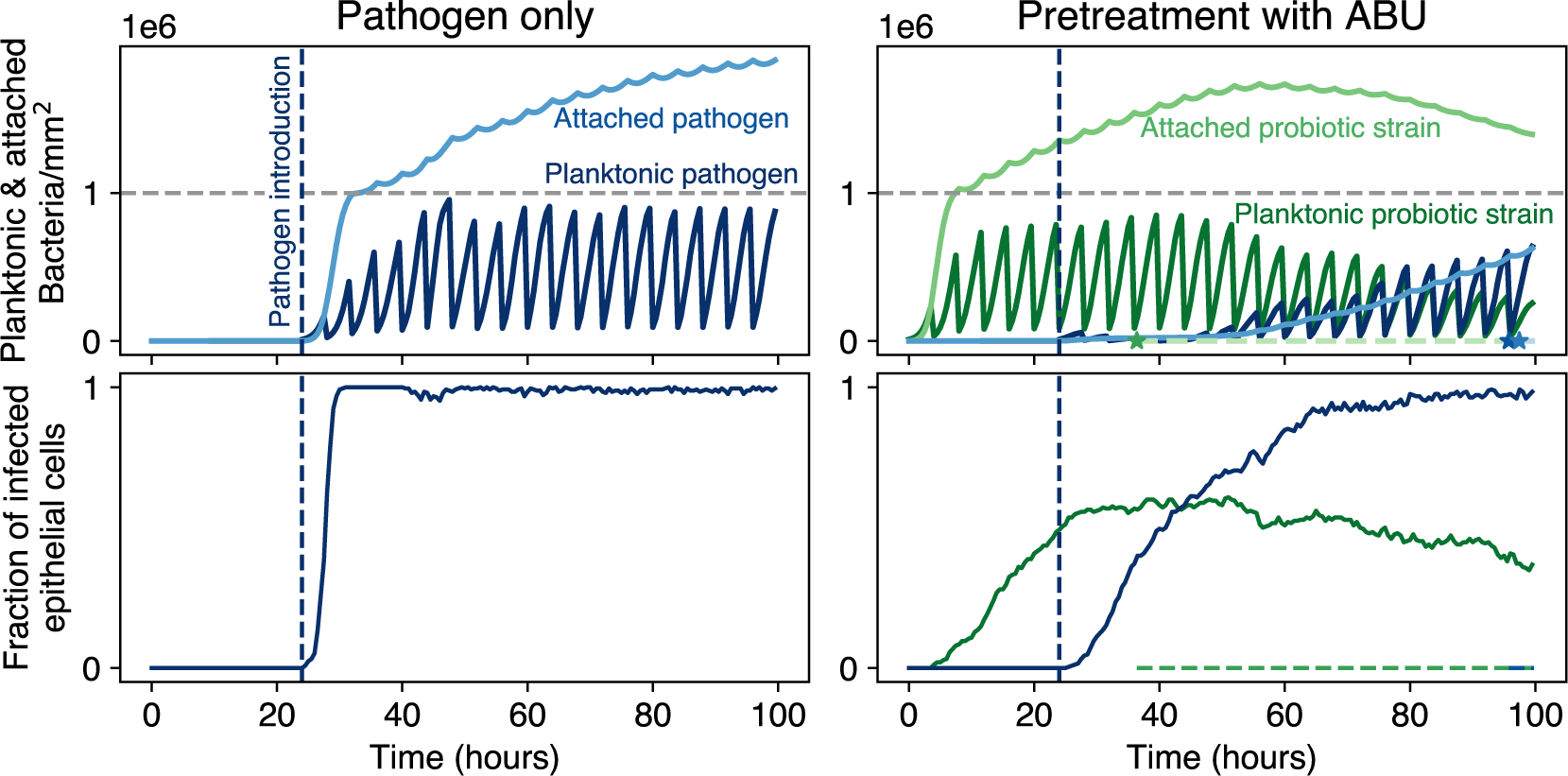
Population dynamics of pathogenic bacteria without (left) or with (right) probiotic pretreatment. The pathogenic bacteria are introduced at *t* = 24 h at a small number (10^4^). For the pretreated case, the system in inoculated with 10^4^ probiotic bacteria in the planktonic compartment at *t* = 0 h. The pathogen has an intermediate invasion rate of *ν*_PATH_ = 10*^−^*^3^ h*^−^*^1^ here, which is two orders of magnitude higher than the invasion rate of the probiotic strain *ν*_ABU_ = 10*^−^*^5^ h*^−^*^1^. The pathogen birth rate is assumed to be 10% below the growth rate of the probiotic strain (*β*_ABU_ = 1.19 h*^−^*^1^, *β*_PATH_ = 1.07 h*^−^*^1^). No antibiotic treatment is applied. The voiding interval is 4 hours. As in Fig. 3, star symbols mark mutation events in the attached compartment, which in this replicate occur only in the pre-treated scenario.

Antibiotic treatment decreases this effect considerably as treatment suppresses also the ABU strain, thus weakening its competitive suppression of the pathogenic strain. However, in combination with antibiotic treatment, probiotic pre-treatment is able to increase the probability of eradicating the pathogen (Fig. 7). In the case of infrequent antibiotic treatment, this increasing success probability is contingent on a considerably lower pathogen birth rate as well as an only slightly increased invasion rate relative to the probiotic strain (Fig. 7b). For frequent treatment, the increase of treatment success probability is more moderate, but independent of pathogen growth rate and invasion rate (Fig. 7c). Probiotic pre-treatment decreases the effective mutation rate of the pathogen (Fig. S5), but may actually increase the total number of mutations arising from both probiotic and pathogenic bacteria due to a higher overall bacteria abundance (Fig. S6). For low pathogen invasion and birth rates, the pretreatment with probiotic bacteria tends to reduce the survival of resistant mutants from the pathogen background, but increases it for weaker trade-offs (i.e. only slight reduction of pathogen birth rate and much higher pathogen invasion rate) (Fig. S7). A similar pattern is apparent when considering the survival of resistant mutants from pathogen and probiotic background together (Fig. S8). These findings indicate the potential of unintentionally aggravating the problem of resistance evolution by prophylactic probiotic pre-treatment.

**Figure 7.**
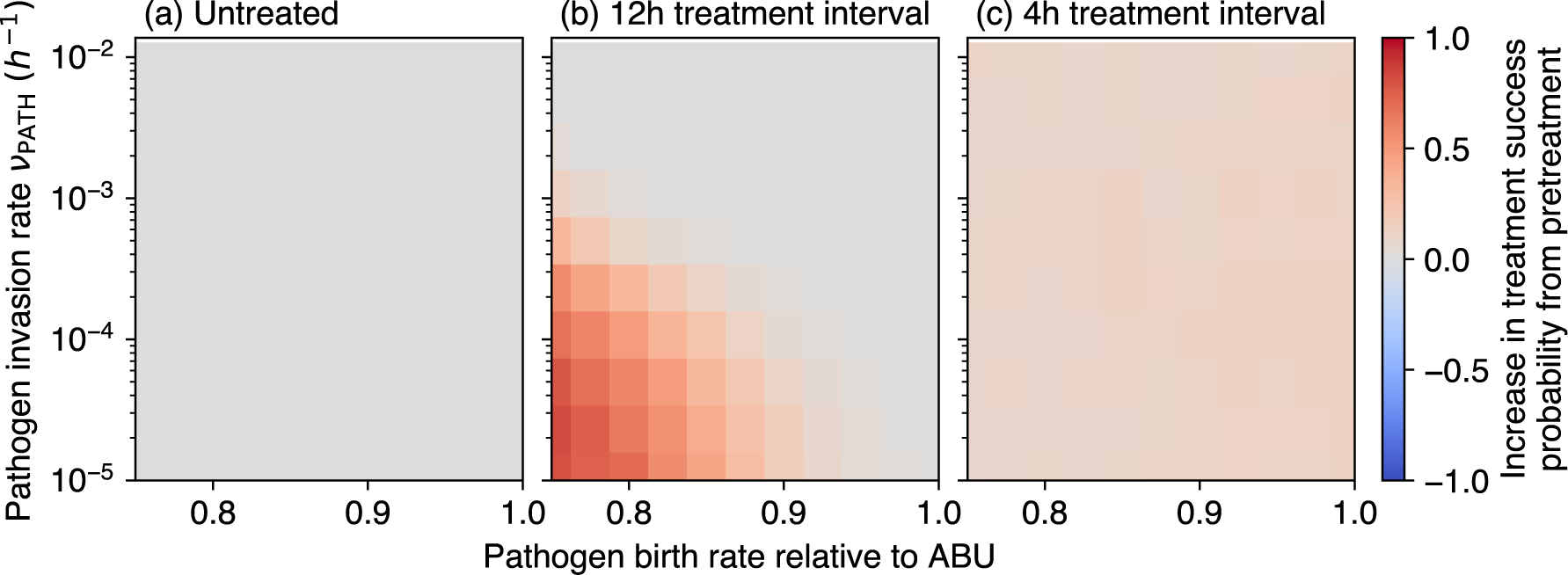
Effect of prophylactic probiotic pretreatment on the extinction probability of the pathogen for different antibiotic perturbation levels. A larger probability of clearing the pathogen is indicated by a positive value (red). As in Fig. 4, pathogen extinction is determined at *t* = 100 h after adding the pathogen at *t* = 24 h. Treatment here is fully permeating (*ϕ* = 1).

## Discussion

After a prior urinary tract infection by *E. coli*, 24% of women suffer a recurrent urinary tract infection within the next six months (Foxman et al., 2000). Besides reinfection from external sources or the intestinal reservoir, persistence within the urinary tract is thought to be a major source of these recurrent infections (Thänert et al., 2019). The intracellular colonization of the bladder epithelium by intracellular bacterial communities (IBCs) is suspected as one explanation for this persistence (Schwartz et al., 2011; Murray et al., 2021). We have presented here a mathematical model of the initial period of urinary tract infections that captures this intracellular compartment in addition to compartments for planktonic bacteria in the bladder lumen and bacteria attached to the urothelium. We have shown that access to the non-planktonic compartments, the invasion rate into the intracellular compartment, as well as the vulnerability of this potential shelter to antibiotic treatment, i.e. the treatment permeation, are decisive parameters for the eco-evolutionary dynamics of initial urinary tract infections.

The different spatial compartments provide distinct growth and perturbation conditions. The planktonic compartment of bacteria is vulnerable both to bladder voiding and antibiotic treatment. The attached compartment is sheltered against voiding but also susceptible to antibiotic treatment. The intracellular compartment may provide shelter against both perturbation types, unless a well-permeating antibiotic drug is used. Transitioning between these compartments thus allows the bacterial population to moderate the differing selection pressures and can generate intricate ecological and evolutionary infection dynamics, which may explain the difficulty of preventing recurrent UTIs.

We observe a trade-off between the chance of eradicating the infection and the risk of resistance evolution. Stronger treatment, i.e. shorter treatment interval or higher permeation, exerts a stronger perturbation but also a higher selection pressure for resistant mutants, thus increasing their survival probability. This general trade-off is well-known and forms the basis for evolution-informed treatment approaches for bacterial infections and cancer (Gatenby and Brown, 2020; Hansen et al., 2020; Burmeister et al., 2020; Merker et al., 2020). These approaches recommend lower treatment-induced selection pressures to delay treatment failure when resistance is already present or when it would inevitably evolve. Our study shows that spatial heterogeneity and refuge from treatment may further complicate the search for the most efficient balance between treatment success and risk of resistance evolution.

We found that intermediate antibiotic permeation can result in long-term, intracellular bacterial populations, similar to persistent IBCs or the recalcitrating effect of biofilms (Trubenová et al., 2022a). We found that these drug-induced intracellular subpopulations can act as mutational hotspots due to the large population size and the high number of replications. These intermediate values of permeation, together with the high treatment frequency, prevent bursting of the epithelial cells without eradicating the intracellular populations, i.e. they could undermine parts of the natural defense against infections. This indicates that accounting for the spatial distribution of perturbations is important to understand the eco-evolutionary dynamics in urinary tract infections, i.e. the location of resistance mutations and their success probability.

Our results show that wildtype population size is a good predictor for the occurrence of mutations, but only in the planktonic and internalized compartment, as attachment flux and shelter from voiding can create attached population sizes beyond the carrying capacity, resulting in the lack of replication. Furthermore, population size is a bad predictor for the survival probability of mutants. Here, the compartment of origin is decisive. Mutants that arise in the attached compartment are more likely to survive our observation window than planktonic mutants. Survival of intracellular mutants depends on high antibiotic selection pressure, i.e. short treatment intervals and high permeation.

One important consequence of access to the intracellular compartment is that the planktonic population size is a bad indicator for the presence or absence of an ongoing infection. For the combination of voiding and antibiotic treatment we find that planktonic population sizes are small, but the attached and intracellular populations are large. As bacteria counts in urine, represented by our planktonic compartment, are the routine surveillance measure for acute urinary tract infections, this poses the risk of too early discontinuation of antibiotic treatment when the infection is not yet fully cured, elevating the risk of antibiotic misuse and widespread resistance evolution. Our findings support recent advocacy for using other diagnostic measures to detect urinary tract infections, such as molecular diagnostic methods (Sathiananthamoorthy et al., 2019; Szlachta-McGinn et al., 2022), which might be particularly important for patients with chronic lower urinary tract symptoms (Khasriya et al., 2020). We find in our model that prophylactic probiotic pretreatment can hinder the establishment of a pathogenic urinary infection, which is supported by clinical evidence (e.g. Sihra et al., 2018). Also, we see a higher success rate of antibiotic treatment with probiotic pretreatment, which is also supported clinically (Mohseni et al., 2013). For infrequent antibiotic treatment, this positive effect increases with a stronger growth advantage of the probiotic strain over the pathogenic one and depends on smaller invasion benefits of the pathogen. Indeed, probiotic candidate strains are often found to grow faster than pathogenic strains (Roos et al., 2006; Stork et al., 2018). However, the effect of probiotic treatment decreases under frequent antibiotic treatment, which is reasonable given our assumption of the antibiotic treatment also targeting the probiotic strain. Judging whether this interaction is also present in patients requires further clinical trials.

In this study, we have focused on the dynamics of early urinary tract infections. Important aspects at later stages include cell-based immune responses, changes of bacterial phenotypes, e.g. filamentation or biofilm formation in the attached phase, and the invasion into deeper layers of the bladder tissue (Flores-Mireles et al., 2015). Although these aspects are certainly important to understand the course of the disease, our focus on the initial stages of UTIs aims at supporting the early resolution and prevention of this frequent infection and thus may contribute to improved health and quality of life for millions of patients.

## Acknowledgements

This research is supported by Dioscuri, a program initiated by the Max Planck Society, jointly managed with the National Science Centre in Poland, and mutually funded by Polish Ministry of Science and Higher Education and German Federal Ministry of Education and Research, grant UMO-2019/02/H/NZ6/00003. M.R. and A.T. are partly supported by the DFG through Research Training Group “Translational Evolutionary Research” (TransEvo) (Project number 400993799). BW is supported by NAWA Polish Returns grant no. PPN/PPO/2019/1/00030/U/0001.

## Competing Interests

The authors declare no competing interests.

## Data availability

All python scripts used in this study have been deposited on Zenodo at https://doi.org/10.5281/ zenodo.10025283. The simulation outputs can be found at https://doi.org/10.5281/zenodo.10025291.

## Supplement

## Supplement

#### Algorithm 1 Pseudo-code for the model describing the initial dynamics of urinary tract infections. Parameter descriptions and their values can be found in Tab. 1.

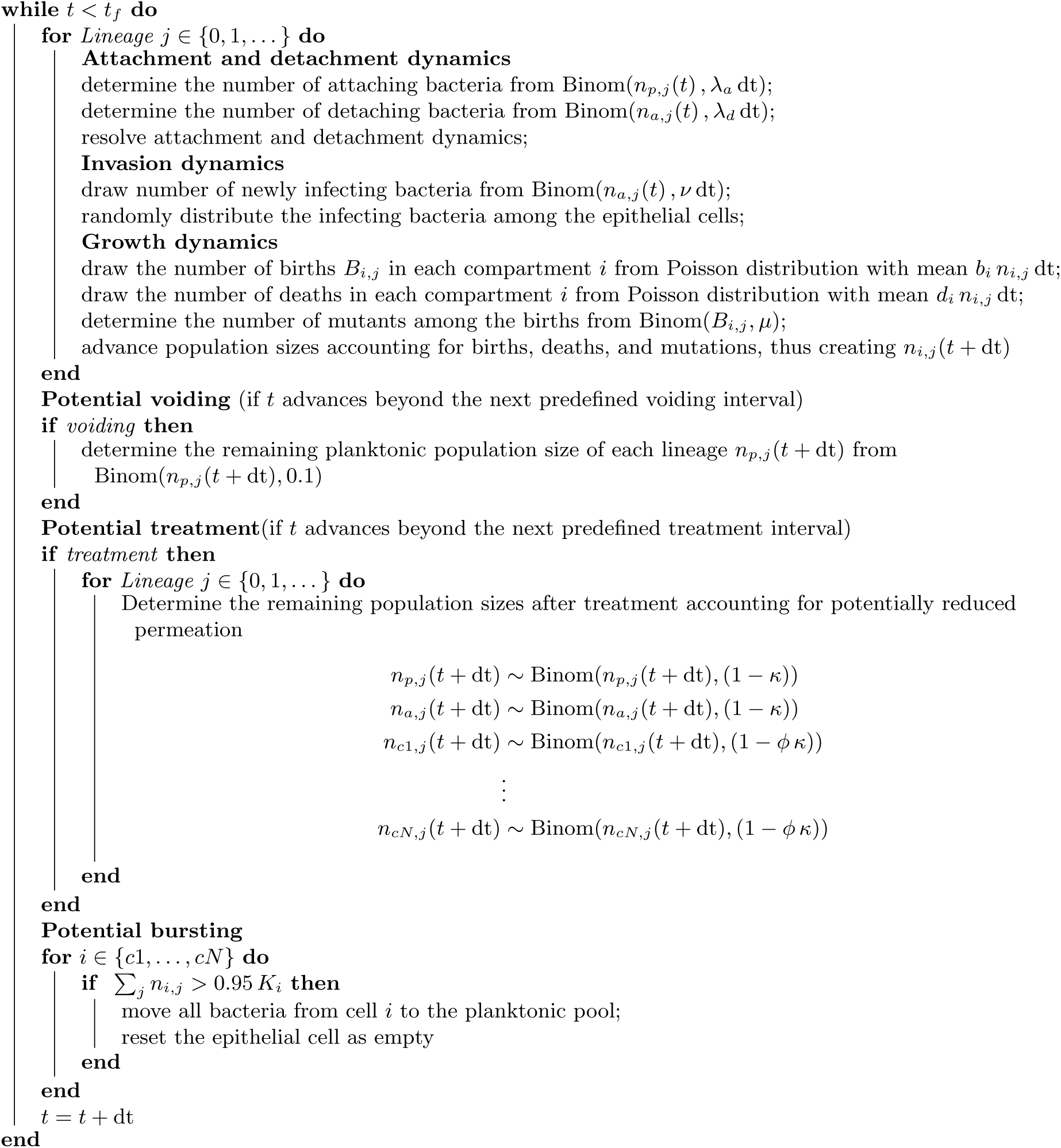

**Figure S1.**
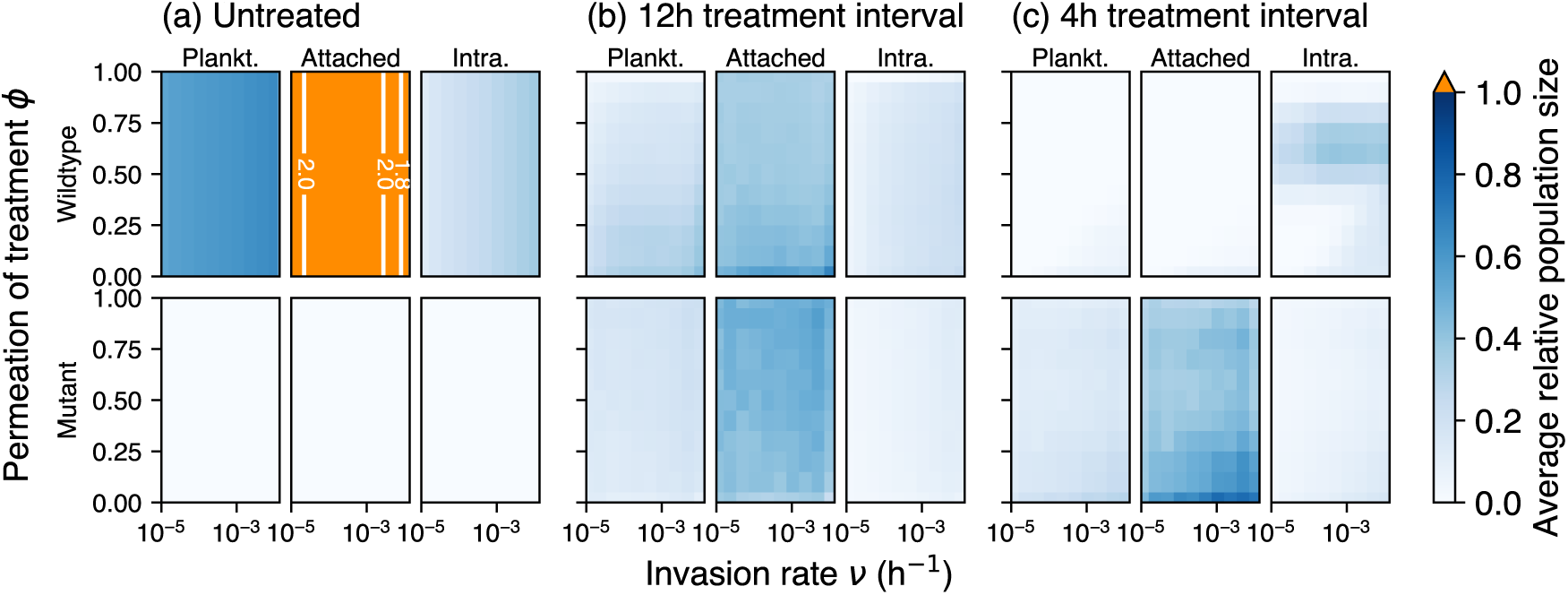
Average relative population size in each of the three compartments. The average population size of wildtype (top row) and mutant (bottom row) across 1000 replicates is plotted relative to the carrying capacity of the respective compartment, i.e. 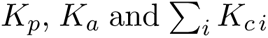 for planktonic, attached and intracellular, respectively. Simulation characteristics as described in Fig. 4. Note that the orange colouring indicates population sizes beyond the carrying capacity of the respective compartment with the white lines giving isoclines for orientation.

**Figure S2.**
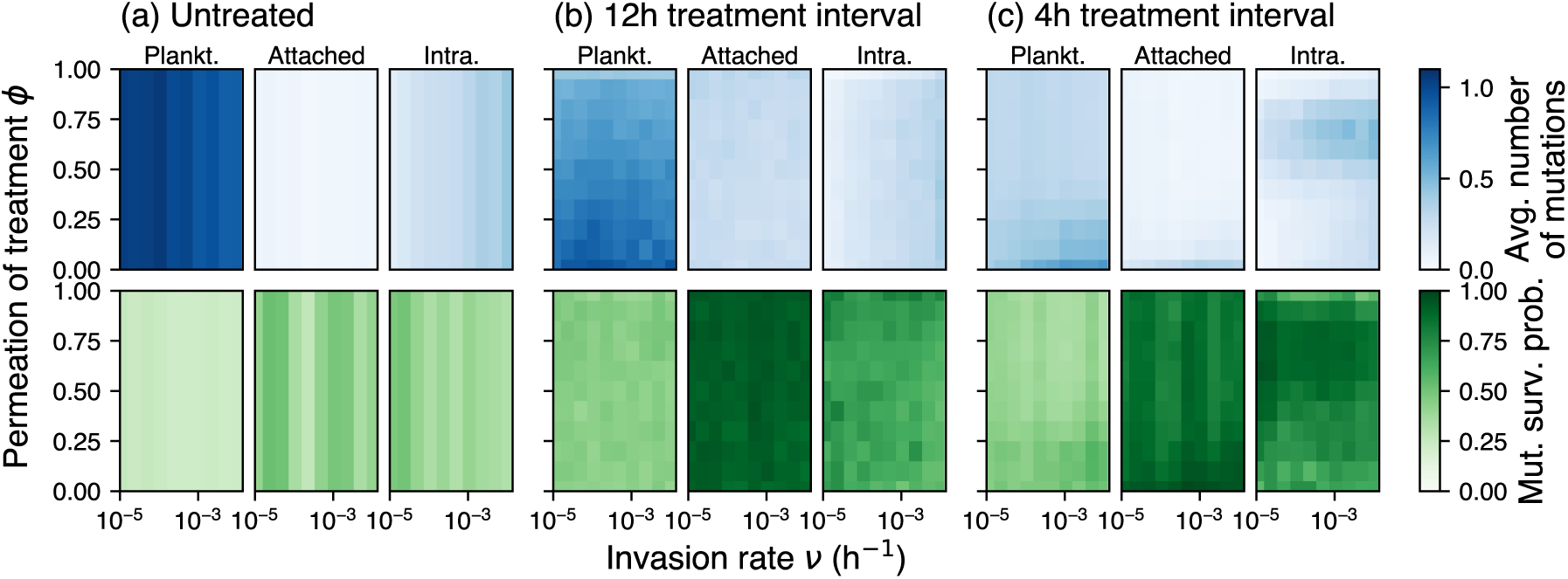
The observed number of mutations and survival probability of mutants for different treatment types in each of the three compartments. Top row: Observed average number of mutations in the respective compartment across 1000 replicates during 100h with treatment start at *t* = 30 h. Bottom row: Probability of survival for these mutants during the observation period. Simulation characteristics as described in Fig. 4.

**Figure S3.**
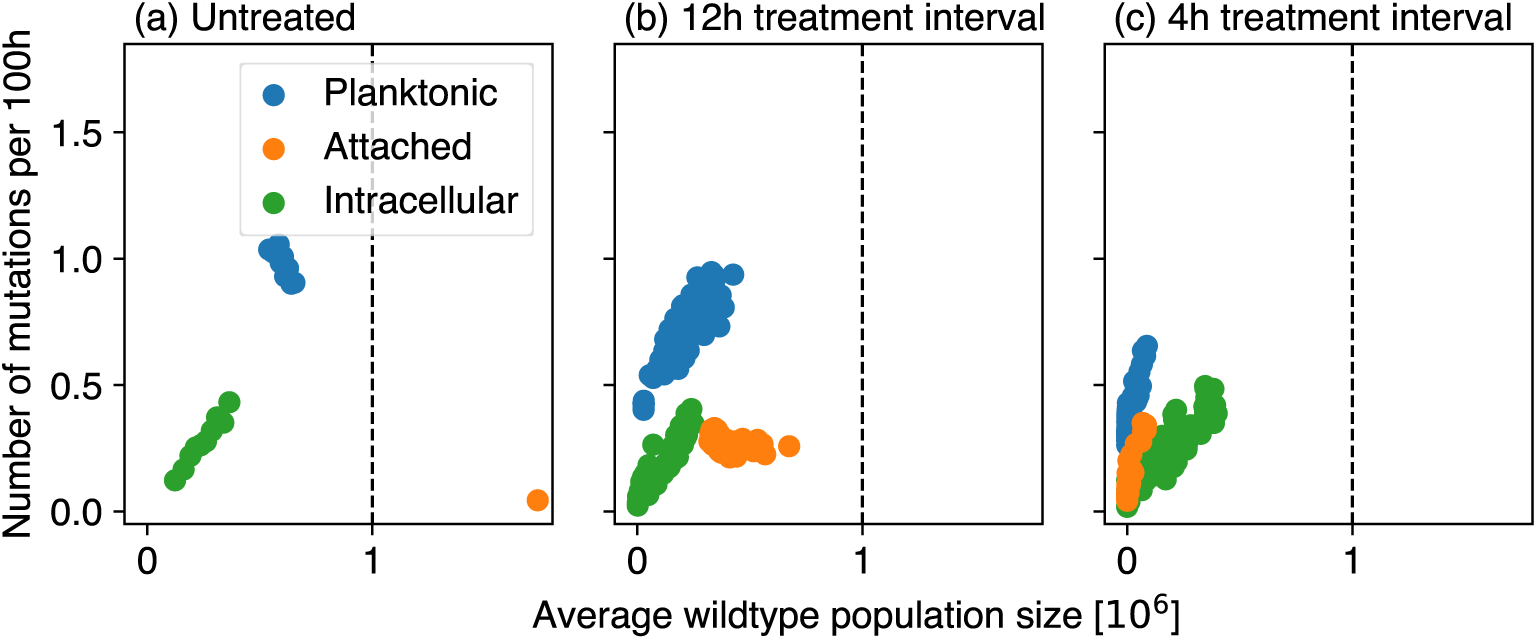
Correlation of observed mutation rate and average wildtype population size in the different antibiotic perturbation scenarios. The points correspond to the different combinations of invasion rate *ν* and treatment permeation *ϕ* explored for example in Fig. 4. Colors indicate the different compartments. The dashed vertical line shows the carrying capacities for birth (assumed to be equal in all compartments). For the intracellular compartment *K_c_* is shown.

**Figure S4.**
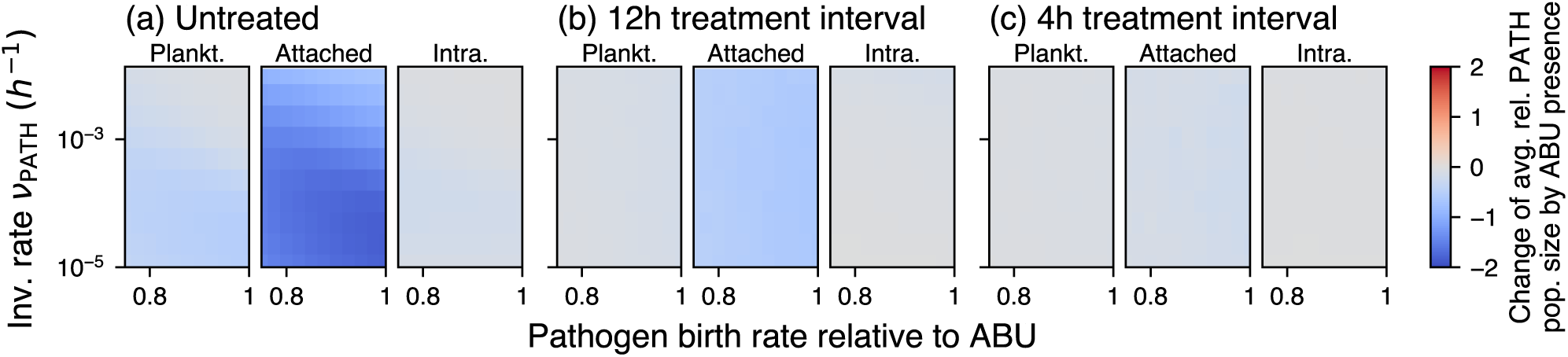
Effect of prophylactic probiotic pretreatment on pathogen population size for different antibiotic treatment intervals. The color gradient shows the change in the relative pathogen population size by compartment averaged over 1000 replicates between *t* = 80h and *t* = 100h. Negative values (blue) indicate a lower average pathogen population size when probiotic pretreatment is added at *t* = 0 h, compared to no probiotic pretreatment. As in Fig. 6, the pathogen is added at 24 h.

**Figure S5.**
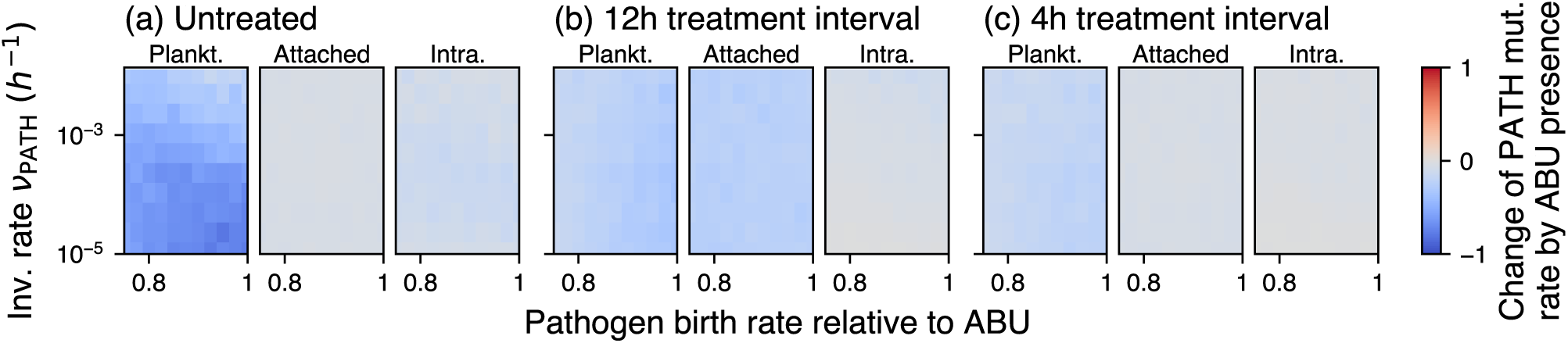
Effect of prophylactic probiotic pretreatment on the observed pathogen mutation rate. Negative values indicate a lower mutation rate on the pathogen background if pretreated with a probiotic strain.

**Figure S6.**
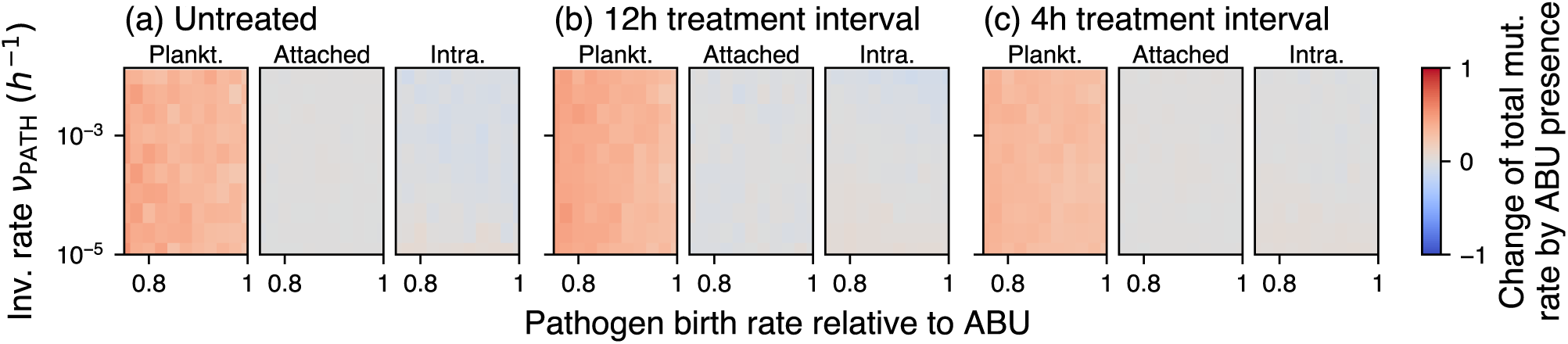
Effect of prophylactic probiotic pretreatment on the observed total mutation rate (including mutations on ABU and PATH background). In contrast to Fig. S5, all mutations are counted here.

**Figure S7.**
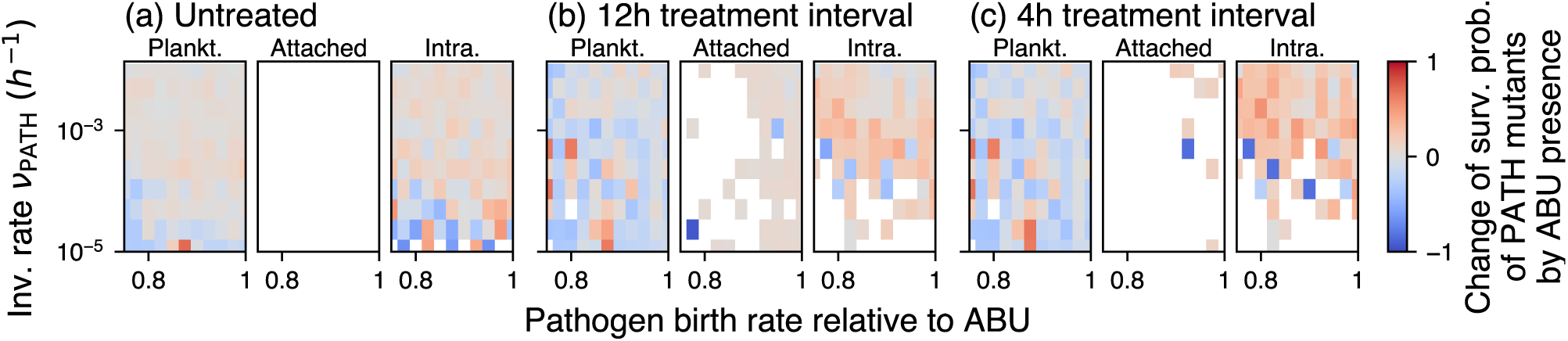
Effect of prophylactic probiotic pretreatment on the survival probability of pathogen mutants. Negative values indicate a lower establishment probability of pathogenic mutants because of prophylactic probiotic pretreatment. Blank values indicate parameter combinations for which mutants did not appear in both scenarios (with and without ABU).

**Figure S8.**
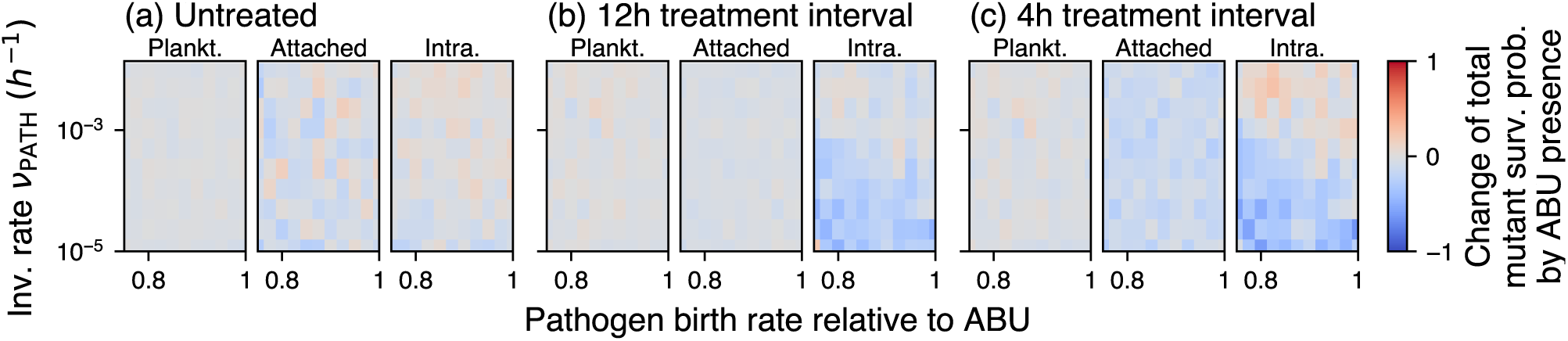
Effect of prophylactic probiotic pretreatment on the survival of either probiotic or pathogen mutants. Negative values indicate a lower establishment probability of mutants from all backgrounds because of prophylactic probiotic pretreatment.

## Notes

https://doi.org/10.5281/zenodo.10025283

https://doi.org/10.5281/zenodo.10025291

### Competing Interest Statement

The authors have declared no competing interest.

